# Polyfunctional IL-21^+^ IFNγ^+^ T follicular helper cells contribute to checkpoint inhibitor diabetes mellitus and can be targeted by JAK inhibitor therapy

**DOI:** 10.1101/2024.11.27.625710

**Authors:** Nicole Huang, Jessica Ortega, Kyleigh Kimbrell, Joah Lee, Lauren N. Scott, Esther M. Peluso, Sarah J. Wang, Ellie Kao, Kristy Kim, Jarod Olay, Zoe Quandt, Trevor E. Angell, Maureen A. Su, Melissa G. Lechner

**Affiliations:** Division of Endocrinology, Diabetes, and Metabolism, University of California Los Angeles (UCLA) David Geffen School of Medicine, Los Angeles, CA 90095; UCSF Medical School, San Francisco, CA 94143; University of Kansas Medical School, Kansas City, KS 66160; UCLA/California Institute of Technology Medical Scientist Training Program, UCLA David Geffen School of Medicine, Los Angeles, CA 90095; California State Polytechnic University, Pomona, CA 91768; Department of Microbiology, Immunology, and Molecular Genetics, UCLA David Geffen School of Medicine, Los Angeles, CA 90095; Division of Endocrinology and Metabolism, University of California San Francisco Medical School, San Francisco, CA 94143; Division of Endocrinology and Diabetes, University of Southern California Keck School of Medicine; Los Angeles, CA 90033; Division of Pediatric Endocrinology, UCLA David Geffen School of Medicine; Los Angeles, CA 90095

## Abstract

Immune checkpoint inhibitors (ICI) have revolutionized cancer therapy, but their use is limited by the development of autoimmunity in healthy tissues as a side effect of treatment. Such immune-related adverse events (IrAE) contribute to hospitalizations, cancer treatment interruption and even premature death. ICI-induced autoimmune diabetes mellitus (ICI-T1DM) is a life-threatening IrAE that presents with rapid pancreatic beta-islet cell destruction leading to hyperglycemia and life-long insulin dependence. While prior reports have focused on CD8^+^ T cells, the role for CD4^+^ T cells in ICI-T1DM is less understood. Here, we identify expansion CD4^+^ T follicular helper (Tfh) cells expressing interleukin 21 (IL-21) and interferon gamma (IFNγ) as a hallmark of ICI-T1DM. Furthermore, we show that both IL-21 and IFNγ are critical cytokines for autoimmune attack in ICI-T1DM. Because IL-21 and IFNγ both signal through JAK-STAT pathways, we reasoned that JAK inhibitors (JAKi) may protect against ICI-T1DM. Indeed, JAKi provide robust *in vivo* protection against ICI-T1DM in a mouse model that is associated with decreased islet-infiltrating Tfh cells. Moreover, JAKi therapy impaired Tfh cell differentiation in patients with ICI-T1DM. These studies highlight CD4^+^ Tfh cells as underrecognized but critical mediators of ICI-T1DM that may be targeted with JAKi to prevent this grave IrAE.

**VISUAL ABSTRACT:** 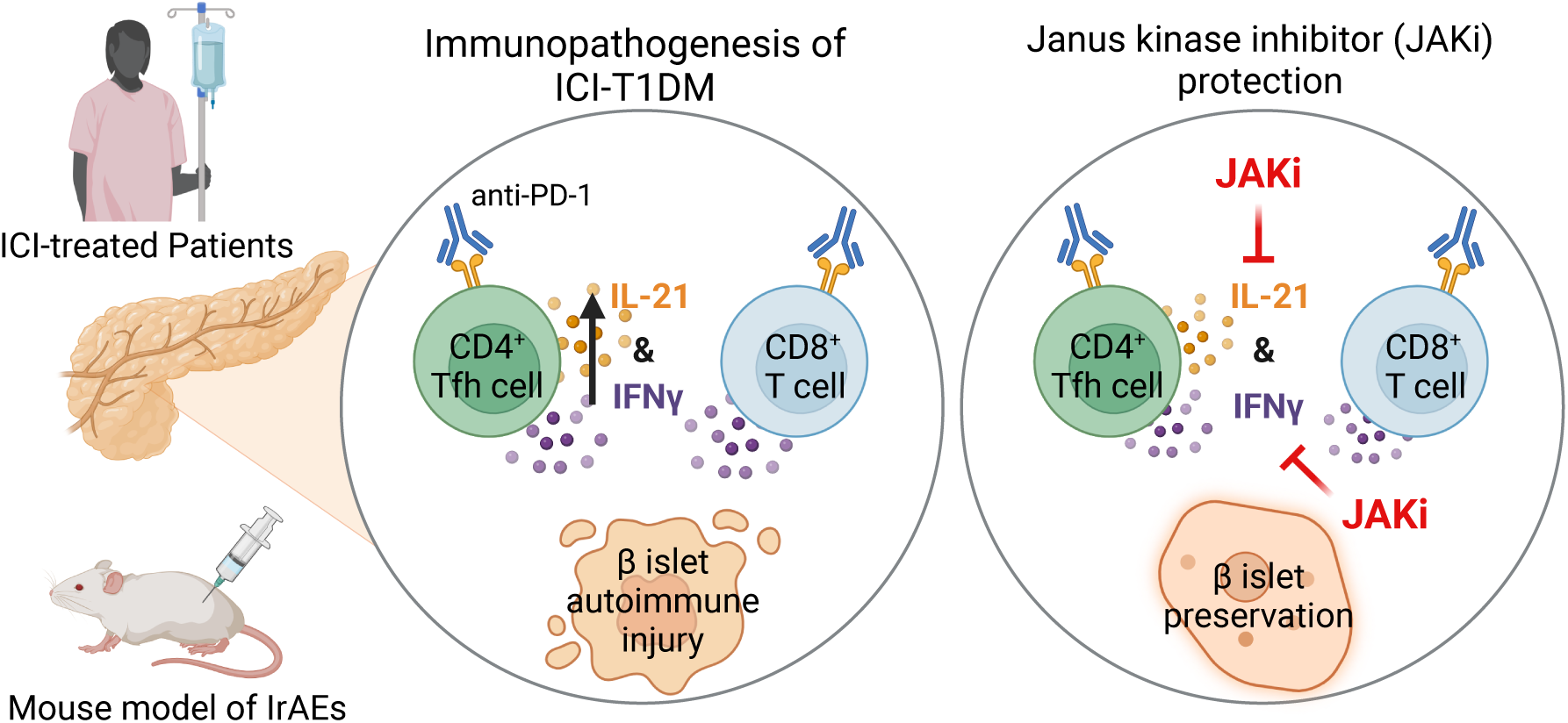

## INTRODUCTION

Immune checkpoint inhibitor (ICI) therapies have significantly improved outcomes for patients with many types of advanced cancers. However, their use is limited by the development of autoimmune toxicities in healthy tissues in nearly two-thirds of patients^1–3^. Autoimmune diabetes mellitus (ICI-T1DM) is a rare but life-threatening immune-related adverse event (IrAE) that occurs in 1-2% of patients treated with ICI^4^. ICI-T1DM presents as a rapidly progressive autoimmune destruction of pancreas beta-islet cells, accompanied by hyperglycemia and often ketoacidosis^4–6^. Patients with ICI-T1DM have permanent pancreatic endocrine insufficiency and require life-long insulin replacement therapy. In patients receiving ICI therapy for advanced malignancies, this additional co-morbidity can add another debilitating and overwhelming layer of complexity to their care. On the other hand, in the growing number of patients who receive ICI therapy for early stage or curable disease, ICI-T1DM represents a permanent sequela of treatment that can negatively impact quality of life long after cancer resolution.

Currently no therapies exist to prevent endocrine IrAEs, including ICI-T1DM^4,5,7–9^. Understanding immune mechanisms that drive autoimmunity may identify therapeutic targets to reduce IrAEs. We recently identified interleukin 21 (IL-21)^+^ T follicular helper (Tfh) cells as critical mediators of ICI-thyroiditis^10^, another common endocrine IrAE seen in 15-25% of ICI-treated patients. Like ICI-T1DM, ICI-thyroiditis presents as brisk autoimmune destruction of thyroid gland cells and loss of thyroid function over a period of weeks^9,11^. We found that thyrotoxic IFNγ^+^ CD8^+^ T cells in the thyroid were driven by IL-21 from CD4^+^ Tfh cells and inhibition of IL-21 prevented ICI-thyroiditis^10^. Whether Tfh cells contribute to the development of ICI-T1DM and may be therapeutically targeted to reduce pancreas autoimmunity during ICI therapy has not yet been explored.

In addition to developing mechanism-based therapies for IrAEs, a practical consideration is the urgent need for near-term strategies to reduce autoimmunity in the many patients currently receiving ICI therapy. As clinical indications for ICI therapy expand^12^, the number of patients with IrAEs will surge – as will the need for therapies to halt severe or life-threatening autoimmune toxicities like ICI-T1DM. Janus kinase inhibitors (JAKi) are a class of orally bioavailable medications now widely used to treat spontaneous autoimmune diseases like alopecia, psoriasis, and arthritis^13–15^. These agents block JAK signaling, which is required for many T cell cytokine responses^13^. Indeed, Waibel et al.^16^ reported preservation of beta-islet cell function and decreased insulin requirements in individuals with spontaneous T1DM in a phase 2 trial of JAKi baricitinib. However, the potential of JAKi to halt the rapid and often fulminant autoimmune responses seen in IrAEs has only been explored recently. JAK 1/2 inhibitor ruxolitinib notably improved survival from 3.4% to 60% in a cohort of patients with steroid refractory ICI-myocarditis, another rare but deadly IrAE, when given in combination with CTLA-4 agonist abatacept^17^. Based upon their promise in spontaneous autoimmune diseases and ICI-myocarditis, we hypothesized that JAKi could be utilized to prevent endocrine IrAEs.

In this study, we identify multifunctional CD4^+^ T follicular helper (Tfh) cells expressing IL-21 and interferon gamma (IFNγ) as antigen-specific mediators of autoimmune tissue injury in ICI-T1DM. Furthermore, we show that both IL-21 and IFNγ are critical cytokines in autoimmune attack during ICI-T1DM and that inhibition of these cytokine pathways by JAKi therapy can prevent ICI-T1DM. Moreover, we show that JAKi treatment decreases islet-infiltrating Tfh cells in a mouse model of IrAEs and Tfh cell differentiation in patients with ICI-T1DM. These studies highlight CD4^+^ Tfh cells as underrecognized but critical mediators of ICI-T1DM that may be targeted with JAKi to prevent this life-threatening endocrine IrAE.

## RESULTS

### Individuals with ICI-T1DM have increased T follicular helper cell responses

T follicular helper cells contribute to multiple spontaneous autoimmune diseases, including T1DM^18,19^, where they can signal to B cells in germinal centers and promote pathogenicity of CD8^+^ T cells^10,18–21^. Expansion of Tfh cells has recently been linked to the development of IrAEs in ICI-treated patients. Herati et al.^22^ reported an increase in circulating Tfh cells after influenza vaccination in anti-PD-1 treated patients who went on to develop IrAEs. Furthermore, in individuals with ICI-thyroiditis, IL-21^+^ CD4^+^ Tfh cells are key drivers of thyroid autoimmune attack^10^. Therefore, we hypothesized that Tfh cells may also contribute to the development of ICI-T1DM.

To test this idea, we evaluated Tfh cells (CD4^+^ ICOS^+^ PD-1^hi^ CXCR5^+^) in peripheral blood specimens from patients with ICI-T1DM vs. patients who received ICI therapy but did not develop IrAEs. Because prior work showed that Tfh cell response, but not baseline levels of circulating Tfh cells, was predictive of IrAEs, we compared the magnitude of Tfh cell expansion between groups after Tfh-skewing *ex vivo* ^23^ (**Fig. 1A**). Indeed, patients with ICI-T1DM had a more robust Tfh cell response than those without IrAEs, with increased CD4^+^ ICOS^+^ PD-1^hi^ CXCR5^+^ cells compared to controls without autoimmunity (**Fig. 1B**, p<0.05). These data suggest that individuals with ICI-T1DM have increased CD4^+^ Tfh cell responses compared to individuals who do not develop IrAEs.

**Figure 1.**
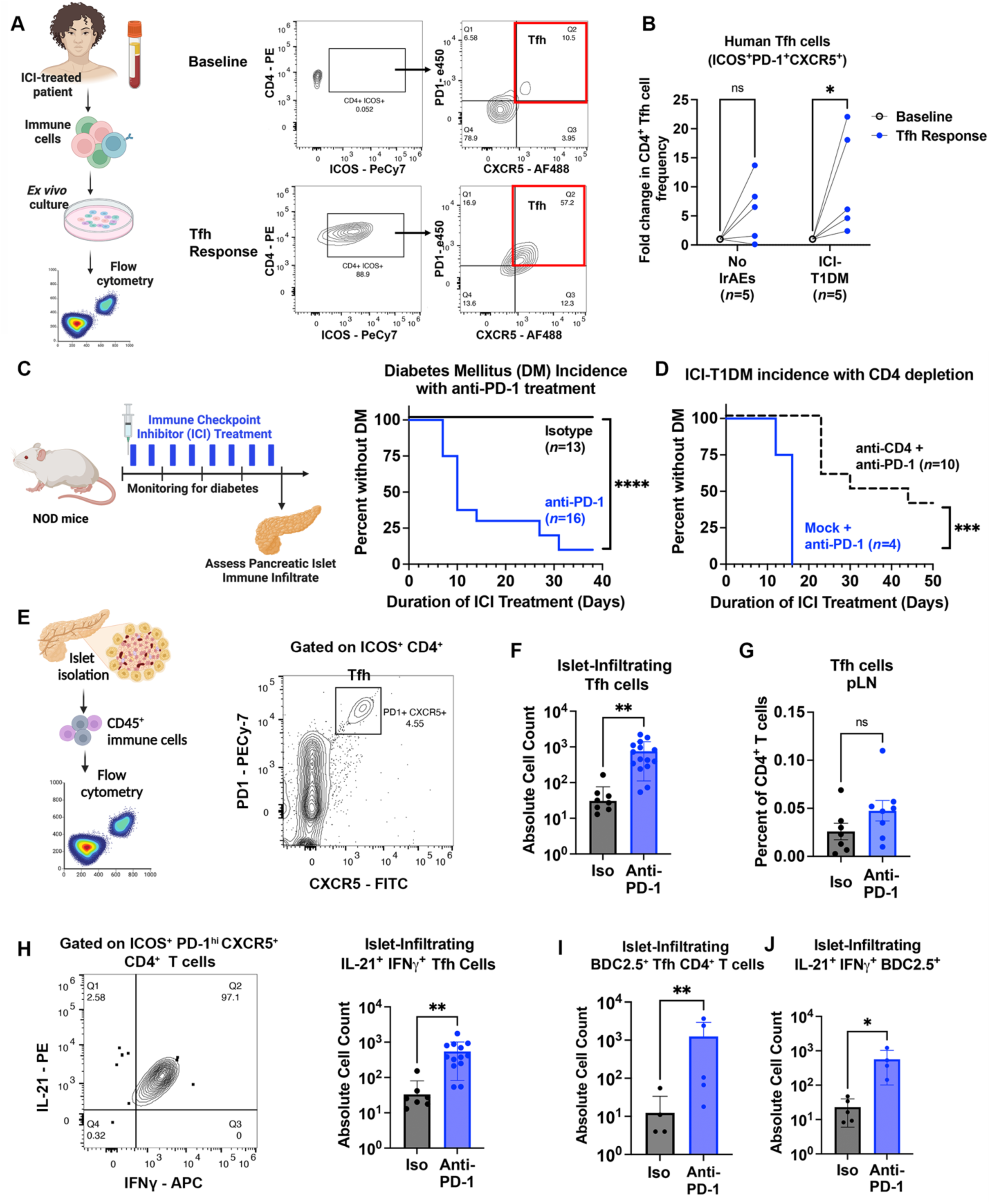
Increased CD4^+^ T follicular helper (Tfh) cell response in individuals with ICI-T1DM and a mouse model of IrAEs. A. Schematic of *ex vivo* Tfh skew of human peripheral blood mononuclear cells from immune checkpoint inhibitor (ICI)-treated cancer patients and assessment of Tfh cells flow cytometry (*left*). Representative flow cytometry plots of Tfh cell markers (CD4, ICOS, PD-1, and CXCR5) at baseline and after *ex vivo* culture under Tfh-skewing conditions for three days as reported previously^23^ (*right*). AF488, Alexa fluorophore emission 488; e450, emission 450 fluorophore; PE, phycoerythrin; PECy-7, phycoerythrin-cyanine 7. B. Comparison of fold change in Tfh cell (CD4^+^ ICOS^+^ PD-1^+^ CXCR5^+^) frequency among individuals with ICI-T1DM and individuals who received ICI-therapy but did not have IrAEs (No IrAEs). Data shown as fold change relative to baseline and each pair represents one individual. C. Schema of IrAE mouse model in which non-obese diabetic (NOD) mice are treated with twice weekly intraperitoneal injections of ICI or isotype control antibodies and monitored for the development of autoimmune diabetes mellitus (DM) (*left*). Incidence of autoimmune DM in NOD mice treated with anti-PD-1 or isotype (Iso). D. DM incidence in anti-PD-1 treated NOD mice with a depleting anti-CD4 antibody or isotype control (Mock). E. Representative flow cytometry plot of CD4^+^ T cells within the pancreatic islets of anti-PD-1 treated mice showing gating for putative T follicular helper (Tfh) cell surface markers. PECy-7, phycoerythrin-cyanine 7; FITC, fluorescein isothiocyanate. F. Quantification of Tfh cells (CD4^+^ ICOS^+^ PD-1^hi^ CXCR5^+^) within the islets of anti-PD-1 compared to Iso treated mice. G. Frequency of Tfh cells within the pancreatic lymph nodes (pLN) of anti-PD-1 compared to Iso treated mice. H. Representative flow cytometry plot showing staining of IL-21 and IFNγ dual cytokine producing CD4^+^ ICOS^+^ PD-1^hi^ CXCR5^+^ Tfh cells in the islet of an anti-PD-1 treated mouse. APC, allophycocyanin; PE, phycoerythrin (*left*). Quantification of islet-infiltrating IL-21^+^ IFNγ^+^ Tfh cells in Iso and anti-PD-1 treated mice (*right*). I. Quantification of BDC2.5^+^ Tfh cells within the islets of Iso versus anti-PD-1 treated mice. J. Comparison of the number of islet-infiltrating IL-21^+^ IFNγ^+^ BDC2.5^+^ CD4^+^ Tfh cells between anti-PD-1 and Iso treated mice. (F-J), Absolute cell counts and frequencies of islet-infiltrating cell types were determined by flow cytometry. Each point represents data from one animal and data are presented as mean±SD. Comparisons by two-way ANOVA for paired samples with subsequent pairwise comparisons (B), Log-Rank test (C,D) or Mann-Whitney test (F-J); *p<0.05, **p<0.01, ***p<0.001,****p<0.0001.

### Antigen-specific, IL-21^+^ IFNγ^+^ CD4^+^ T follicular helper cells are increased in the pancreatic islets of mice with ICI-T1DM

To better understand the role of Tfh cells in the immunopathogenesis of ICI-T1DM *in vivo*, we then used a mouse model of IrAEs. Previously, we reported the development of multi-organ immune infiltrates in autoimmunity-prone non-obese diabetic (NOD) mice following ICI treatment, including thyroiditis, colitis, and accelerated diabetes mellitus^10,24^. As expected, male and female NOD mice (7-9 weeks of age) treated with continued cycles of anti-programed death protein (PD-1) antibody (10mg/kg/dose, twice weekly), developed ICI-T1DM at a median of 10 days, while isotype treated controls remained healthy after four weeks (**Fig. 1C**, p<0.0001).

T cells play a key role in the development of IrAEs in multiple tissues^10,25–28^, including the pancreas^4,29–31^. As expected, NOD mice with genetic deletion of the TCRα gene, which leads to an absence of mature CD4^+^ and CD8^+^ T cells, were completely protected from ICI-T1DM (**Suppl. Fig. 1A**). Prior studies have demonstrated the importance of IFNγ-producing CD8^+^ T cells in mouse models of ICI-T1DM^29–31^. On the other hand, the role of CD4^+^ T cells has been less explored but is important in other IrAEs^10,24,32–34^. Additionally, because CD4^+^ T cell responses may not be as central to ICI anti-tumor efficacy, they might be potential therapeutic targets to reduce IrAEs in patients with cancer while preserving efficacy^24,32,35^.

Antibody depletion of CD4^+^ T cells in ICI-treated wildtype NOD mice significantly delayed the onset of autoimmune diabetes (**Fig. 1D**, p<0.001 and **Suppl. Fig. 1B**), suggesting a CD4^+^ T cell contribution to ICI-T1DM disease progression. We then compared the frequency of CD4^+^ Tfh cells within pancreatic islets and pancreatic lymph nodes (pLN) of NOD mice after three weeks of anti-PD1 or isotype control therapy (**Fig. 1E**). Indeed, anti-PD1 treated mice had increased islet-infiltrating Tfh cells (CD4^+^ ICOS^+^ PD-1^hi^ CXCR5^+^) compared to isotype controls (**Fig. 1F**, p<0.01); a trend toward increased Tfh cells was also found in pLN, but this difference was not statistically significant (**Fig. 1G**). We next evaluated cytokine production of islet-infiltrating Tfh cells by flow cytometry and found an increase in dual producing IL-21^+^ IFNγ^+^ Tfh cells in anti-PD-1 treated mice (**Fig. 1H**, p<0.01). Such multifunctional IL-21^+^ IFNγ^+^ Tfh CD4^+^ cells have previously been described as mediators of immune response in spontaneous autoimmune diseases (e.g. lupus and peripheral neuropathy) and viral infections ^36–39^.

Fife and colleagues previously established a pathogenic role for BDC2.5^+^ CD4^+^ T cells in NOD mice with accelerated autoimmune DM due to loss of PD-1^40^. Therefore, we used an MHC class II BDC2.5 tetramer to quantify auto-antigen-specific CD4^+^ T cells in our mouse model. Twenty-seven percent of islet-infiltrating BDC2.5^+^ CD4^+^ T cells had a surface phenotype consistent with Tfh cells (ICOS^+^ PD-1^hi^ CXCR5^+^) by flow cytometry (**Suppl. Fig. 2A** and **2B**), expressed canonical Tfh transcription factor b cell lymphoma 6 (Bcl6)^+^ (**Suppl. Fig 2C** and **2D**), and produced cytokines IL-21 and IFNγ (**Suppl. Fig. 2E** and **2F**). Furthermore, anti-PD-1 treated mice had more BDC2.5^+^ CD4^+^ Tfh cells in pancreatic islets compared to isotype-treated controls (**Fig. 1I**, p<0.01) and these cells showed high dual expression of IL-21 and IFNγ (**Fig. 1J**, p<0.05). Taken together, these data support a role for antigen-specific, polyfunctional IL-21^+^ IFNγ^+^ CD4^+^ Tfh cells in the autoimmune attack on pancreas beta-islet cells during ICI therapy.

### IL-21 and IFNγ are important cytokine mediators of ICI-T1DM

We hypothesized that inhibition of Tfh cytokines, specifically IL-21 and IFNγ (**Fig. 2A**), could attenuate autoimmune attack on the pancreas during anti-PD-1 therapy. IL-21 is a pleiotropic cytokine that can promote effector functions in CD8^+^ T cells ^10,20,21^ and B cell antibody production^41^. In humans and mice, CD4^+^ Tfh cells are the primary source of IL-21^18,42^. Indeed, NOD mice with genetic deletion of IL-21 signaling (NOD.IL21RKO) were protected from the development of ICI-T1DM during ICI treatment (**Fig. 2B**, p<0.0001 for anti-PD-1 therapy in WT versus IL21RKO mice). It is recognized that IL-21 is required for the development of spontaneous T1DM in NOD mice^43,44^, and these data establish a role for IL-21 in ICI-T1DM as well.

**Figure 2.**
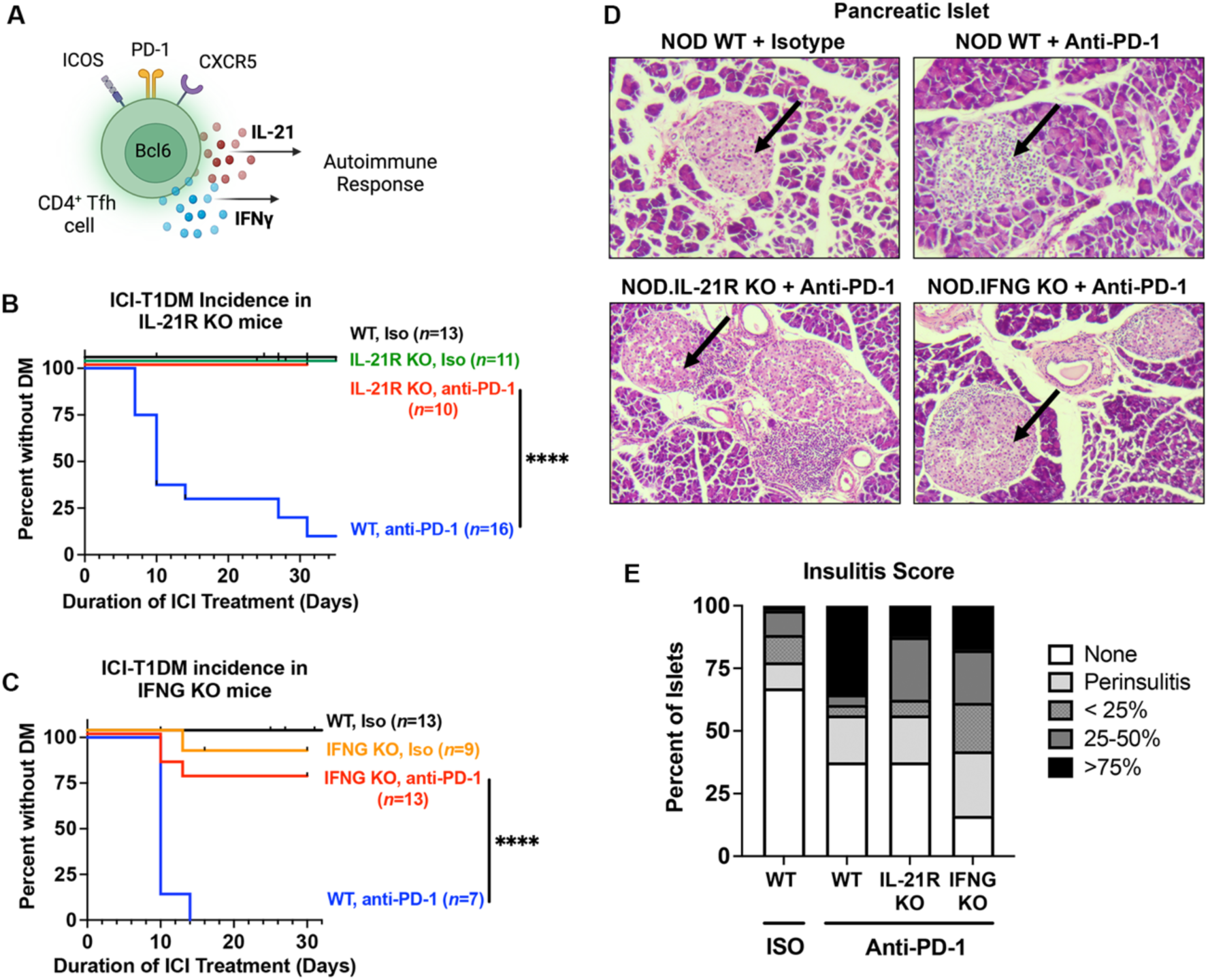
Interleukin 21 (IL-21) and interferon gamma (IFNγ) are key cytokine mediators of ICI-T1DM. A. Schematic of cytokine production by Tfh cells. B. Incidence curve for ICI-T1DM in anti-PD-1 treated NOD. WT and NOD.IL-21R KO mice. C. Incidence curve for ICI-T1DM in ICI-treated NOD.WT and NOD.IFNG KO mice during anti-PD-1 treatment. D. Representative hematoxylin and eosin-stained pancreas histology sections of isotype (Iso) or anti-PD-1 treated NOD WT, IL-21R KO, or IFNG KO mice (original magnification 100X). Arrow indicates an islet of Langerhans. E. Insulitis index determined by histologic analyses of pancreas islet histology across indicated treatment conditions. Comparisons by Log-Rank test (B, C). ****p<0.0001.

IFNγ is expressed more broadly, including by both CD4^+^ and CD8^+^ T cells in ICI-T1DM^29–31^. NOD mice with genetic deletion of the IFNγ gene (NOD.IFNG KO) showed significantly delayed onset of ICI-T1DM (**Fig. 2C**, p<0.0001 for anti-PD-1 therapy in WT versus IFNG KO mice). These data confirm a previous non-significant trend reported by Perdigoto et al.^31^ and are consistent with IFNγ as a mediator of ICI-T1DM^29^. Pancreas histology and insulitis scoring of both anti-PD-1-treated NOD.IL21R KO and NOD.IFNG KO mice confirmed reduced islet infiltration compared to anti-PD-1 treated WT mice (**Fig. 2D and 2E**). In summary, our data identify a role for IL-21^+^ IFNγ^+^ Tfh cells in ICI-T1DM and demonstrate that inhibition of these two cytokine pathways can prevent the development of autoimmunity.

### JAK1/2 inhibition via ruxolitinib prevents ICI-induced diabetes mellitus in NOD mice

With the expanding use of ICI therapies and the rising number of patients affected by IrAEs, there is a pressing clinical need for near-term strategies to prevent or reverse treatment-associated autoimmunity. To this end, we wondered whether JAK inhibitors, a group of clinically-approved agents used in spontaneous autoimmune diseases^45–48^, could prevent ICI-T1DM. JAK signaling is central to many T cell immune responses, including downstream signals of IL-21^41^ and IFNγ^49^ (JAK1/2 and JAK1/3, respectively) (**Fig. 3A**). Using our mouse model, we tested whether treatment with JAK1/2 inhibitor ruxolitinib could delay development of ICI-T1DM (**Fig. 3B**). Notably, while anti-PD-1 treated mice on control food rapidly developed autoimmune diabetes, ruxolitinib therapy prevented ICI-associated autoimmunity, with no mice developing overt DM (**Fig. 3B**, p<0.0001).

**Figure 3.**
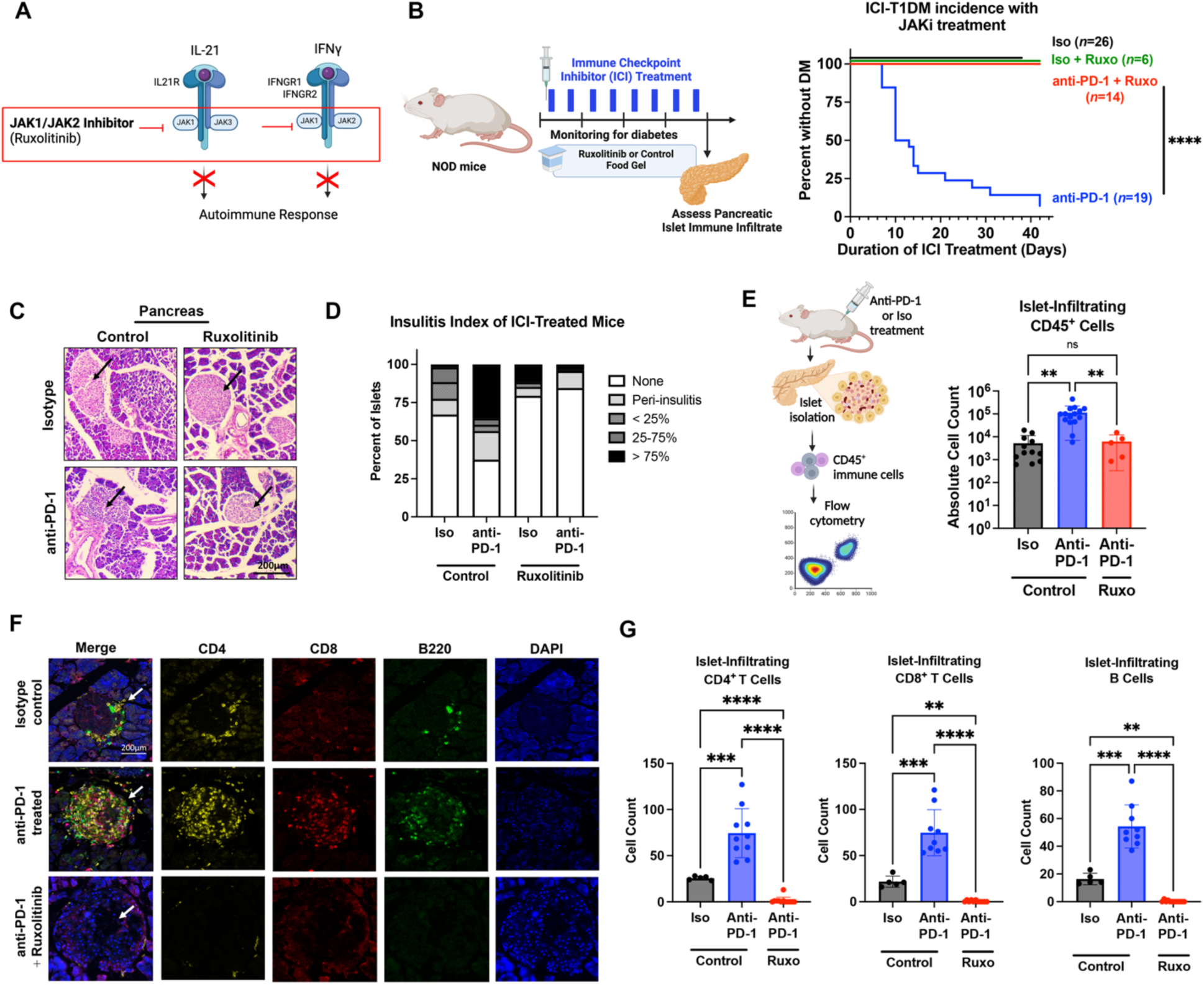
Janus kinase inhibitor (JAKi) ruxolitinib provides robust protection against immune checkpoint inhibitor (ICI) autoimmune diabetes mellitus (DM). A. Proposed targeting of JAK signaling mediating downstream from IL-21 and IFNγ to halt autoimmune response. treatment of mice with JAKi ruxolitinib to reduce ICI-T1DM. B. Schematic for treatment of mice with JAKi ruxolitinib (*left*) and incidence of autoimmune DM (*right*) in NOD mice treated with anti-PD-1 immunotherapy or isotype (Iso) and ruxolitinib or control food gel. C. Representative hematoxylin and eosin-stained pancreas histology sections of anti-PD-1 or Iso-treated NOD mice (original magnification 100X) fed ruxolitinib or control food. Arrow indicates an islet of Langerhans. D. Insulitis index of anti-PD-1 or Iso-treated NOD mice given ruxolitinib or control food. E. Schematic and absolute cell counts of pancreatic islet-infiltrating CD45^+^ cells, as determined by flow cytometry, across Iso + vehicle (*n*=12), anti-PD-1 + vehicle (*n*=15), and anti-PD-1 + ruxolitinib (*n*=5) conditions. Each point represents data from one animal. F. Representative multi-immunofluorescence staining and microscopy images (original magnification 40X) of CD4, CD8, B220, and DAPI in the islet of Langerhans across experimental conditions. Arrow indicates islet in merge images. G. Quantification of CD4^+^ T cell, CD8^+^ T cell, and B220^+^ B cell counts per pancreatic islet of indicated treatment condition by immunofluorescence. Data are presented as mean±SD (E,G). Comparisons by Log-Rank test (B) or ANOVA with Welch’s correction and pairwise comparison by Tukey’s test (E, G). **p<0.01, ***p<0.001, ****p<0.0001.

Histologic analysis of pancreatic islets from anti-PD-1-treated mice given ruxolitinib showed minimal immune infiltrate (**Fig. 3C**), with insulitis scores comparable to isotype-treated controls (**Fig. 3D**). We confirmed reduced immune infiltrates in ruxolitinib-fed mice using flow cytometry analysis of immune cells in isolated pancreatic islets. Compared to anti-PD-1 treated mice, those additionally given ruxolitinib had significantly reduced islet-infiltrating CD45^+^ immune cells (p<0.01), comparable to isotype-treated controls (p=ns) (**Fig. 3E**). Our further characterization and quantification of immune infiltrates in pancreatic islet infiltrates using multi-parameter immunofluorescence staining of tissue specimens (**Fig. 3F**) demonstrated that CD4^+^ T cells, CD8^+^ T cells, and B220^+^ B cells accumulated within the islets of anti-PD-1 treated mice and were significantly decreased by the addition of ruxolitinib therapy (**Fig. 3G**, p<0.0001 for all cell types).

In addition, we tested whether ruxolitinib could reverse ICI-T1DM. This is clinically relevant because the development of ICI-T1DM in patients is usually detected after clinical diabetes development. Here, groups of anti-PD-1 treated mice were randomized to treatment with ruxolitinib or vehicle control after the development of diabetes (blood glucose >200mg/dL) (**Suppl. Fig. 3A**). Ruxolitinib treated mice had improved glycemic control compared to vehicle controls (**Suppl. Fig. 3B**, p<0.0005) and had decreased insulitis scores on pancreas histology (**Suppl. Fig. 3C**). Finally, prior studies have shown PD-1 blockade to be the primary driver of accelerated autoimmunity in adult mice^50–52^, but ICI-T1DM can also develop in cancer patients treated with combination ICI regimens [e.g. anti-PD-1 + anti-cytotoxic T lymphocyte antigen (CTLA-4)]^4–6^. As in mice that received anti-PD-1 monotherapy, JAKi treatment prevented the development of ICI-T1DM in mice treated with combination anti-PD-1 + anti-CTLA-4 (**Suppl. Fig. 4**). In summary, these data show potent *in vivo* protection against ICI-T1DM using a clinically available JAKi.

### JAKi therapy disrupts the CD4^+^ Tfh cell compartment to prevent ICI-T1DM

IL-21 signaling to CD4^+^ T cells supports Tfh cell differentiation and relies upon JAK signaling. As such, we predicted that JAKi therapy may attenuate Tfh cell responses by preventing autocrine IL-21 signaling in CD4^+^ T cells (**Fig. 4A**). Indeed, CD4^+^ T cells with genetic IL-21 receptor loss (IL-21R KO) had reduced differentiation to Tfh cells *in vitro* (**Fig. 4B**, p<0.05).

**Figure 4.**
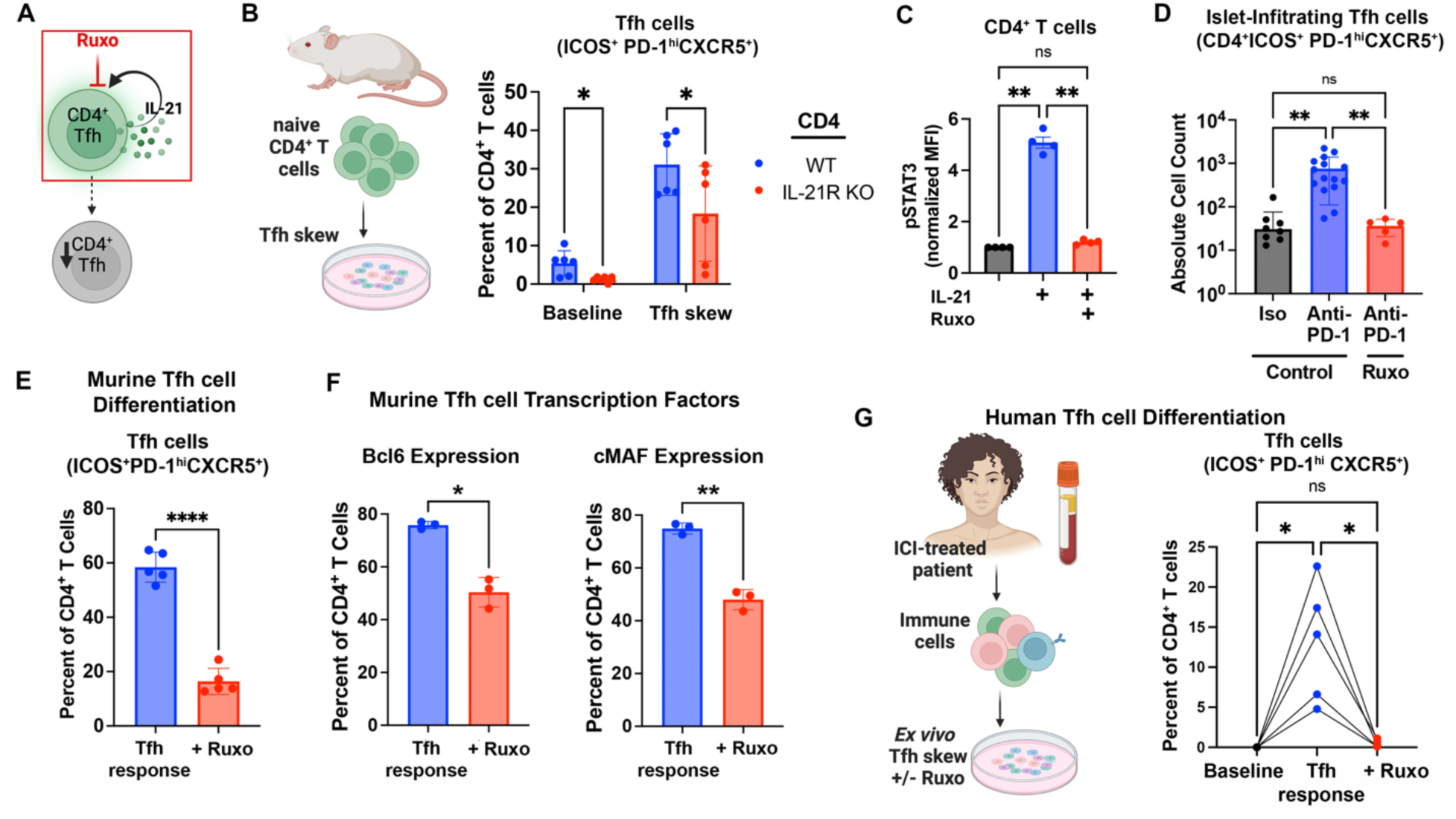
JAKi treatment reduces CD4^+^ Tfh cell response in mice and humans. A. Schematic of proposed action of JAKi on CD4^+^ Tfh cells through blockade of autocrine IL-21 downstream signaling. B. Impact of IL-21 receptor genetic deletion in CD4^+^ T cells on the induction of Tfh cells from naïve CD4^+^ T cells *in vitro*, assessed after three days under Tfh-skew conditions with anti-PD-1. C. Phosphorylation of STAT3 in murine CD4^+^ T cells in response to IL-21 (100ng/mL) ruxolitinib (10uM) or vehicle control *in vitro*, assessed by flow cytometry. D. Quantification of pancreatic islet-infiltrating, IL-21^+^ IFNγ^+^ co-producing Tfh cells as determined by flow cytometric analysis, among Iso + vehicle (n=7), anti-PD-1 + vehicle (n=13), and anti-PD-1 + ruxolitinib (n=5) treated mice. E. Frequency of murine CD4^+^ T cells expressing a Tfh cell phenotype (CD4^+^ ICOS^+^ PD-1^hi^ CXCR5^+^) following a three-day Tfh skew of naïve CD4^+^ T cells in the presence of ruxolitinib (10uM) or vehicle control *in vitro*, assessed by flow cytometric analysis. F. Expression of canonical Tfh transcription factors Bcl6 and cMAF in murine CD4^+^ T cells following a three day Tfh skew in the presence ruxolitinib (Ruxo, 10uM) or vehicle control. G. Comparison of Tfh cell response in PBMC specimens from ICI-treated individuals at baseline and following a three-day Tfh skew with JAKi ruxolitinib (Ruxo, 10uM) or vehicle control, measured by flow cytometry. Each point represents data from one replicate (B, C, E, F; experiments repeated at least twice) or animal (D), and data are presented as mean±SD. For human studies (G), connected points represent data from one individual. Comparisons by two-way ANOVA (B) or one-way ANOVA (C, D, G) with subsequent pairwise comparisons or Welch’s t test (E, F). *p<0.05; **p<0.01, ****p<0.0001.

Additionally, *in vitro* treatment of CD4^+^ T cells with ruxolitinib prevented JAK-mediated intracellular STAT3 phosphorylation in response to IL-21 stimulation (**Fig. 4C**, p<0.01). Moreover, JAKi treatment in our mouse model of ICI-T1DM led to significantly fewer CD4^+^ Tfh cells (ICOS^+^ PD-1^hi^ CXCR5^+^) within pancreatic islets (**Fig. 4D**, p<0.01). These data show that in addition to blocking the downstream effects of Tfh cell cytokines (i.e. IL-21 and IFNγ), JAKi therapy is associated with decreased Tfh cells *in vivo*.

To better understand the impact of JAKi on Tfh cell expansion in ICI-treated mice, we evaluated naïve CD4^+^ T cells under Tfh skewing conditions^53^ with and without ruxolitinib *in vitro.* Flow cytometry analysis revealed that ruxolitinib reduced markers of Tfh cell differentiation in CD4^+^ T cells (**Fig. 4E**, p<0.0001) and down regulated the expression of Bcl6 (p<0.05) and cMAF (p<0.01), a transcription factor required for IL-21 expression in Tfh cells (**Fig. 4F**).

We then evaluated whether JAKi treatment similarly impaired Tfh cell responses in humans. Using peripheral blood specimens from patients treated with ICI therapy, we compared the frequency of CD4^+^ Tfh cells after culture under Tfh-skewing conditions *ex vivo*. Indeed, Tfh cell differentiation was significantly decreased by JAKi ruxolitinib (**Fig. 4G**, p<0.05). Thus, JAKi decreases Tfh cell induction in murine and human CD4^+^ T cells, suggesting a mechanism by which the development of ICI-T1DM can be prevented *in vivo*. Taken together, these data support JAK inhibitors as a potential near-term therapeutic strategy by which we can target CD4^+^ Tfh cell responses in patients with ICI-T1DM.

## DISCUSSION

The benefits of immune checkpoint inhibitor therapy hold great promise for patients with many types of cancer, but their use is limited by autoimmune adverse events. Among the most severe IrAEs is ICI-induced diabetes mellitus (ICI-T1DM), which leads to destruction of pancreatic islets and life-long insulin dependence. In addition, because of the rapid progression of islet loss compared to spontaneous autoimmune T1DM, patients with ICI-T1DM more frequently present with diabetic ketoacidosis (nearly 80-90%) at diagnosis and require hospital admission to an intensive care unit^6,54^.

In this study, we provide evidence for robust protection of ICI-T1DM with ruxolitinib, an FDA-approved and clinically available JAK1/2 inhibitor. This builds upon a prior report by Ge at al.^55^ evaluating a pre-clinical selective JAK 1 inhibitor for ICI-T1DM in mice and the recent successful phase 2 trial of JAKi baricitinib in spontaneous T1DM^16^. Furthermore, two studies combining JAKi and anti-PD1 therapy in patients with non-small cell lung cancer or Hodgkin’s lymphoma reported improved cancer outcomes^56,57^. Thus, JAKi therapies may be a feasible and near-term approach to reducing toxicity from severe IrAEs such as ICI-T1DM.

On the other hand, JAK signaling is important for many T cell immune responses and can induce broad immune suppression when used spontaneous autoimmune diseases. It is possible, therefore, that JAKi treatment may also impair desired immune responses during cancer immunotherapy. Thus, one aim of our present study was to delineate the cellular mechanisms underlying immune protection during JAKi therapy so that more targeted immunosuppressive strategies could be developed. Prior studies have shown that effector IFNγ^+^ CD8^+^ T cells contribute to autoimmune attack on pancreatic beta-islet cells during ICI therapy^29–31^. Given the importance of IFNγ and CD8^+^ T cells to ICI anti-tumor immune responses^35^, we focused on the less explored role of CD4^+^ T cells with the aim of identifying driving immune mechanisms that could be targeted in cancer patients to reduce IrAEs while preserving efficacy.

Indeed, we found a significant contribution from CD4^+^ T cells in the immunopathogenesis of ICI-T1DM. Specifically, our findings support a critical role for CD4^+^ T follicular helper cells in the immunopathogenesis of ICI-T1DM. Within islet immune infiltrates, we demonstrated antigen-specific multifunctional IL-21^+^ IFNγ^+^ CD4^+^ Tfh cells that were enriched in ICI-treated mice with diabetes. Wherry and colleagues also showed expansion of circulating CD4^+^ Tfh cells in anti-PD-1 treated patients after influenza vaccination correlated with the development of IrAEs (p=0.06)^22^. Expanding upon this role in the periphery, we showed increased Tfh cells in the thyroid tissue of patients with ICI-thyroiditis^10^ and now extend their role to ICI-T1DM. Our data also highlight Tfh cells as a source of two important cytokines in the autoimmune response, namely IFNγ and IL-21. Hu and colleagues previously showed the IFNγ contributes to immune cell migration into pancreas islets and macrophage activation in ICI-T1DM^29^. While a role for IL-21 in ICI-T1DM has not been described, we showed in mice and humans with ICI-thyroiditis that IL-21 from CD4^+^ cells could augment effector molecules (IFNγ, granzyme B) and chemokine receptors on thyrotoxic CD8^+^ T cells^10^. In addition, IL-21 is well known to promote spontaneous T1DM^43^.

To further explore how JAKi may modulate this pathogenic CD4^+^ T cell subset, we evaluated Tfh cells in both ICI-treated mice and *ex vivo* using human specimens. In both contexts, we observed decreased Tfh cell frequency. While JAKi are known to block the downstream effects of multiple T cell cytokines^41,45^, we showed that they can also block Tfh cell differentiation.

These dual mechanisms may be collectively responsible for the benefit of JAKi treatment seen in spontaneous and cancer immunotherapy-associated autoimmune diseases. Furthermore, our studies revealed a vulnerability in autocrine IL-21 CD4^+^ T cell signaling as a potential targeted approach to reduce ICI-associated autoimmunity. Future studies are warranted to further evaluate JAK inhibition and the IL-21 CD4^+^ Tfh cell axis as potential therapeutic targets for reversal of severe IrAEs in ICI-treated patients, as well as the impact on ICI anti-tumor immune responses. In conclusion, our studies not only indicate strong pre-clinical application of JAK1/2 inhibition in the protection of ICI-T1DM development but demonstrate a critical role for IL-21^+^ INFγ^+^ CD4^+^ Tfh cells in driving the mechanism of autoimmune attack and pancreatic injury in ICI-T1DM.

## METHODS

### Sex as a biologic variable

IrAEs occur in both males and females. Therefore, for animal studies, both male and female mice were used in equal proportions. For human studies, both male and female subjects were eligible for participation and included.

### Antibodies and Reagents

Primary immune cells were cultured in RPMI-1640 complete media [supplemented with 10% fetal bovine serum (FBS), 2mM L-glutamine, 1mM HEPES, non-essential amino acids, and antibiotics (penicillin and streptomycin)], with 50µM beta-mercaptoethanol (2ME) (all reagents from Thermo Fisher). Immune checkpoint inhibitor antibodies used were anti-mouse PD-1 (clone RPM1-14, BE0146), CTLA-4 (clone 9D9, BE0164), and isotype control (clone 2A3 BE0089) (all from BioXcell). Antibodies were diluted in sterile PBS for use. Ruxolitinib was obtained from MCE and diluted in sterile DMSO (Sigma) for *in vitro* use. For animal experiments, ruxolitinib was prepared at 1g/kg in Nutra-Gel Diet (Bio-Serv, F5769-KIT) chow as previously described^58^.

### Mouse studies

Animal studies were approved by the UCLA Animal Research Committee (Protocols C21-039 and C24-012). NOD/ShiLtJ (NOD, #001976), NOD/Il21r^-/-^ (IL-21R KO, #034163), NOD/Trca^-/-^ (TCRα KO, #004444), NOD/IFNγ^-/-^ (IFNG KO, #002575), and NOD/SCID (#001303) mice were obtained from the Jackson Laboratory. Male and female mice were used in equal proportions. Mice were used at 7 to 9 weeks of age unless otherwise noted. Mice were housed in a specific pathogen-free barrier facility at UCLA. Mice in different experimental groups were co-housed.

### Immune checkpoint inhibitor treatment of mice

Mice were randomized to continuous twice-weekly treatment with anti-mouse PD-1 (clone RPM1-14) and/or CTLA-4 (clone 9D9) or isotype control antibody (clone 2A3), at 10 mg/kg/dose intraperitoneally (i.p.), as described previously^24^. During treatment, mice were monitored daily for activity and appearance, and twice weekly for weight and glucosuria. Mice developing glucosuria or blood glucose >200mg/dL were treated with 10 units of subcutaneous NPH insulin daily. At the end of ICI treatment course, mice were euthanized and perfused with 10mL of sterile phosphate-buffered saline (PBS) by intracardiac puncture, and fresh tissues were immediately collected for histology or dissociated for analysis of immune infiltrates by flow cytometry. Predetermined endpoints for early euthanasia included >20% weight loss and glucosuria not resolved by insulin therapy, as per IACUC protocols.

For the evaluation of immune infiltrates by flow, islet-infiltrating lymphocytes were collected following an isolation protocol described by Villarreal et al.^59^. Fresh pancreas specimens were perfused with 3mL of a collagenase P solution (1mg/mL Collagenase P in HBSS, supplemented with 0.05% BSA) via the ampulla of Vater, dissected away from surrounding tissue, and mechanically digested in a 37°C thermo-shaker at 100-120 rpm for 13 minutes. Pancreatic islets were then purified using a Histopaque-1077 density gradient (Sigma-Aldrich). Spleen cells were isolated by mechanical dissociation and passage through a 40µm filter.

### Ruxolitinib treatment of mice

For animal experiments, ruxolitinib was prepared at 1g/kg in Nutra-Gel Diet (Bio-Serv, F5769-KIT) chow as previously described^58^. Nutra-Gel Diet chow without ruxolitinib served as control food. For DM reversal experiments, mice were treated with anti-PD-1 twice weekly and monitored daily for blood glucose levels by tail prick. Mice with hyperglycemia (blood glucose level >200 mg/dL), were randomized into either ruxolitinib or control chow groups and ICI treatment was stopped. Ruxolitinib therapy was given as a single oral gavage dose of 1.25mg on day of hyperglycemia onset, followed by ruxolitinib chow (1g/kg) as above. All diabetic mice were assessed daily for blood glucose and treated with NPH insulin if hyperglycemic (10 units for blood glucose >300mg/dL, 5 units for blood glucose between 200 and 300mg/dL). Mice with persistent hyperglycemia not resolved by insulin therapy after four days were euthanized per IACUC protocol.

### *In vitro* assessment of primary murine immune cells

Splenocytes were isolated from healthy NOD.WT mice by mechanical dissociation. Naïve CD4^+^ T and CD8^+^ T cells were isolated by magnetic bead separation as above and cultured at 5×10^5^ cells/well in 12 well plates in complete media with 2ME. For Tfh skew, cells were stimulated with plate bound anti-murine CD3 (Invitrogen, clone 145-2C11; 1 μg/mL) and soluble anti-murine CD28 (Invitrogen, clone 37.51; 1 μg/mL), anti-murine IFNγ (BioXCell XMG1.2,10 μg/mL), anti-murine IL-4 antibodies (BD Biosciences, catalog 554385; 10 μg/mL), anti–murine TGF-β (Thermo Fisher Scientific, catalog 16-9243-85; 20 μg/mL), recombinant mouse IL-6 (PeproTech; 10 ng/mL) and IL-21 (PeproTech; 10 ng/mL), as previously described^53^. Cells were evaluated on day three by flow cytometry. Experiments were repeated at least twice.

### Histology and Immunofluorescence

Harvested tissues were fixed in Zinc for at least 48 hours and then stored in 70% ethanol. Organs were embedded in paraffin, sectioned (4m), and stained with hematoxylin and eosin (H&E) by the UCLA Translational Pathology Core Laboratory. Insulitis quantified by blinded assessment of H&E sections as previously reported^60^. Images were acquired on an Olympus BX50 microscope using Olympus CellScans Standard software. Images were brightened uniformly for publication in Photoshop.

Antibody clones and dilutions: Appropriate positive and negative controls were used for all stains. DAPI, Opal 520 stain for B Cells, Opal 570 stain for CD4^+^ T cells, and Opal 690 stain for CD8^+^ T cells were used to stain. Images of the islet were exported to Photoshop and then analyzed in ImageJ for total islet area and quantification of each cell type.

### Patients

Peripheral blood specimens from cancer patients treated with immune checkpoint inhibitor therapy (ICI) were collected from patients treated in endocrinology and oncology clinics at UCLA or UCSF under Institutional Review Board-approved protocols. Patients were stratified for development of IrAEs , including ICI-T1DM, during ICI therapy versus those with no history of IrAEs. ICI-T1DM was defined by new onset hyperglycemia with low c-peptide and insulin dependence during ICI cancer therapy, consistent with current guidelines from the National Comprehensive Cancer Network (NCCN) Management of Immunotherapy Toxicities guidelines^7^. Other IrAEs were classified based upon NCCN and Common Terminology Criteria for Adverse Events (CTCAE v5) criteria. No IrAE individuals had no evidence of any grade :22 IrAE or pre-existing autoimmune disease. Individual data are presented in **Suppl. Table 1**.

### *In vitro* assessment of primary human immune cells

Peripheral blood mononuclear cells were isolated from whole blood by density gradient centrifugation using Ficoll-Paque (Cytiva). For Tfh skew, cells were stimulated with plate bound anti-human CD3 (BioXcell, clone UCHT1; 1 ug/mL) and soluble anti-human CD28 (BioXcell, clone 9.3; 1 ug/mL), recombinant human IL-12 (PeproTech; 10 ng/mL) and Activin A (R&D Systems; 100 ng/mL), as previously described^23^. Cells were evaluated on day three by flow cytometry. Experiments were repeated at least twice.

### Flow Cytometry

For staining, single-cell suspensions were resuspended in FACS buffer consisting of 0.5mM EDTA, and 2% FBS in phosphate-buffered saline (PBS) at 10^6^ cells/mL. Cells were stained in LIVE/DEAD Fixable Yellow Dead Cell Stain (ThermoFisher) for 30 minutes prior to surface staining. Cells were then stained with fluorescence-conjugated antibodies as indicated in **Suppl. Table 2**. For intracellular staining, after surface staining, cells were fixed and permeabilized using cytoplasmic fixation and permeabilization kit (BD Biosciences), per manufacturer instructions, with a 20-minute fixation step at 4°C. To assess intracellular cytokines, cells were incubated in complete RPMI-1640 media with 50µM 2ME for 4 hours with ionomycin (1µg/mL) (Thermo Fisher) and PMA (50ng/mL) (Sigma) in the presence of Brefeldin A (Biolegend) before staining. For intranuclear staining of phosphorylated signaling proteins, cells were fixed and permeabilized with 4% paraformaldehyde (PFA) for 15 minutes at room temperature, and ice-cold 100% methanol at 4°C for 45 minutes. For intranuclear staining of transcription factors, cells were fixed and permeabilized using a transcription factor fixation and permeabilization buffer kit (Thermo Fisher), following the provided manufacturer protocol including a 30-minute fixation at room temperature. For tetramer staining, BDC2.5 mimotope (CD4^+^ T cell) fluorescently conjugated reagents were obtained from the NIH Tetramer Core and stained at room temperature for 30 minutes as previously described^61^. After staining, cells were washed twice in FACS buffer and analyzed by flow cytometry on an Attune NxT 6 cytometer (Thermo Fisher).

Cell counts are shown as the relative frequency of live, gated single cells unless otherwise noted. For the determination of infiltrating cells within pancreatic islets, absolute cell counts were determined using counting beads (Thermo Fisher, C36995), following the manufacturer’s protocol. Beads were added to pancreatic islet samples at a concentration of 1uL/7uL of sample volume. Representative gating strategies are shown in **Figure 1** and **Suppl. Fig. 1** and **2**.

### Statistical analysis

Statistical analyses were performed using GraphPad Prism software (v10). Comparisons among multiple groups for continuous data were made using ANOVA or ANOVA with Welch correction with no assumption for equal variances, with subsequent pairwise comparisons by Tukey’s or Dunnett’s test. Non-parametric data were evaluated using the Mann-Whitney test.

Comparisons between two groups were done by two-sided Student’s t-test with Welch correction with no assumption for equal variances. Differences in diabetes incidence over time were compared using Log Rank test. When multiple comparisons were performed, adjusted p-values were shown. Significance was defined as α = 0.05.

### Study approval

All animal experiments were conducted under UCLA IACUC-approved protocols and complied with the Animal Welfare Act and the National Institutes of Health guidelines for the ethical care and use of animals in biomedical research. All human experiments were conducted under UCLA and UCSF IRB-approved protocols.

### Data availability

Data comprising figures is provided in Supporting Data Values file. Additional data available upon reasonable request.

## Acknowledgments

This work was supported by grants from the National Institutes of Health (K08 DK129829, M.G.L.), the Doris Duke Charitable Foundation (M.G.L.), and the Aramont Charitable Foundation (M.G.L.). Patient studies were facilitated by the Parker Institute for Cancer Immunotherapy at UCLA. Flow cytometry was performed in the UCLA Jonsson Comprehensive Cancer Center (JCCC) that is supported by National Institutes of Health awards P30 CA016042. We thank the NIH Tetramer Core Facility (contract number 75N93020D00005) for providing BDC2.5 tetramers.

## References Cited

1 Wei SC, Duffy CR, Allison JP. Fundamental mechanisms of immune checkpoint blockade therapy. Cancer Discov 2018;8:1069–86. 10.1158/2159-8290.CD-18-0367.

2 Molina GE, Zubiri L, Cohen J V., Durbin SM, Petrillo L, Allen IM, et al. Temporal Trends and Outcomes Among Patients Admitted for Immune-Related Adverse Events: A Single-Center Retrospective Cohort Study from 2011 to 2018. Oncologist 2021;26:514–22. 10.1002/onco.13740.

3 Wang DY, Salem JE, Cohen J V., Chandra S, Menzer C, Ye F, et al. Fatal Toxic Effects Associated With Immune Checkpoint Inhibitors: A Systematic Review and Meta-analysis. JAMA Oncol 2018;4:1721–8. 10.1001/jamaoncol.2018.3923.

4 Bluestone JA, Anderson M, Herold KC, Stamatouli AM, Quandt Z, Perdigoto AL, et al. Collateral Damage: Insulin-Dependent Diabetes Induced with Checkpoint Inhibitors.

5 Kotwal A, Haddox C, Block M, Kudva YC. Immune checkpoint inhibitors: An emerging cause of insulin-dependent diabetes. BMJ Open Diabetes Res Care 2019;7:. 10.1136/bmjdrc-2018-000591.

6 Wright JJ, Salem JE, Johnson DB, Lebrun-Vignes B, Stamatouli A, Thomas JW, et al. Increased reporting of immune checkpoint inhibitor-associated diabetes. Diabetes Care 2018:e150–1. 10.2337/dc18-1465.

7 Thompson JA, Schneider BJ, Brahmer J, Andrews S, Armand P, Bhatia S, et al. NCCN Guidelines Insights: Management of Immunotherapy-Related Toxicities, Version 1.2023. J Natl Compr Canc Netw 2023;18:230–41. 10.6004/jnccn.2020.0012.

8 Faje AT, Lawrence D, Flaherty K, Freedman C, Fadden R, Rubin K, et al. High-dose glucocorticoids for the treatment of ipilimumab-induced hypophysitis is associated with reduced survival in patients with melanoma. Cancer 2018;124:3706–14. 10.1002/cncr.31629.

9 Ma C, Hodi FS, Giobbie-Hurder A, Wang X, Zhou J, Zhang A, et al. The impact of high-dose glucocorticoids on the outcome of immune-checkpoint inhibitor–related thyroid disorders. Cancer Immunol Res 2019;7:1214–20. 10.1158/2326-6066.CIR-18-0613.

10 Lechner MG, Zhou Z, Hoang AT, Huang N, Ortega J, Scott LN, et al. Clonally-Expanded, Thyrotoxic Autoimmune Mediator CD8+ T cells Driven by IL21 Contribute to Checkpoint Inhibitor Thyroiditis. Sci Transl Med 2023;15:eadg0675. 10.1126/scitranslmed.adg0675.

11 Iyer PC, Cabanillas ME, Waguespack SG, Hu MI, Thosani S, Lavis VR, et al. Immune-Related Thyroiditis with Immune Checkpoint Inhibitors. Thyroid 2018;28:1243–51. 10.1089/thy.2018.0116.

12 Haslam A, Prasad V. Estimation of the Percentage of US Patients With Cancer Who Are Eligible for and Respond to Checkpoint Inhibitor Immunotherapy Drugs. JAMA Netw Open 2019;2:e192535. 10.1001/jamanetworkopen.2019.2535.

13 Tanaka Y, Luo Y, O’Shea JJ, Nakayamada S. Janus kinase-targeting therapies in rheumatology: a mechanisms-based approach. Nat Rev Rheumatol 2022;18:133–45. 10.1038/s41584-021-00726-8.

14 Słuczanowska-Głąbowska S, Ziegler-Krawczyk A, Szumilas K, Pawlik A. Role of Janus Kinase Inhibitors in Therapy of Psoriasis. J Clin Med 2021;10:. 10.3390/jcm10194307.

15 King BA, Craiglow BG. Janus kinase inhibitors for alopecia areata. J Am Acad Dermatol 2023;89:S29–32. 10.1016/j.jaad.2023.05.049.

16 Waibel M, Wentworth JM, So M, Couper JJ, Cameron FJ, MacIsaac RJ, et al. Baricitinib and β-Cell Function in Patients with New-Onset Type 1 Diabetes. New England Journal of Medicine 2023;389:2140–50. 10.1056/NEJMoa2306691.

17 Salem JE, Bretagne M, Abbar B, Leonard-Louis S, Ederhy S, Redheuil A, et al. Abatacept/Ruxolitinib and Screening for Concomitant Respiratory Muscle Failure to Mitigate Fatality of Immune-Checkpoint Inhibitor Myocarditis. Cancer Discov 2023;13:1100–15. 10.1158/2159-8290.CD-22-1180.

18 Crotty S. Follicular helper CD4 T cells (TFH). Annu Rev Immunol 2011;29:621–63. 10.1146/annurev-immunol-031210-101400.

19 Walker LSK. The link between circulating follicular helper T cells and autoimmunity. Nat Rev Immunol 2022;22:567–75. 10.1038/s41577-022-00693-5.

20 Niogret J, Berger H, Rebe C, Mary R, Ballot E, Truntzer C, et al. Follicular helper-T cells restore CD8 + -dependent antitumor immunity and anti-PD-L1/PD-1 efficacy. J Immunother Cancer 2021;9:. 10.1136/jitc-2020-002157.

21 Zander R, Kasmani MY, Chen Y, Topchyan P, Shen J, Zheng S, et al. Tfh-cell-derived interleukin 21 sustains effector CD8+ T cell responses during chronic viral infection. Immunity 2022;55:475–493.e5. 10.1016/j.immuni.2022.01.018.

22 Herati RS, Knorr DA, Vella LA, Silva LV, Chilukuri L, Apostolidis SA, et al. PD-1 directed immunotherapy alters Tfh and humoral immune responses to seasonal influenza vaccine. Nat Immunol 2022;23:1183–92. 10.1038/s41590-022-01274-3.

23 Locci M, Wu JE, Arumemi F, Mikulski Z, Dahlberg C, Miller AT, et al. Activin A programs the differentiation of human T FH cells. Nat Immunol 2016;17:976–84. 10.1038/ni.3494.

24 Lechner MG, Cheng MI, Patel AY, Hoang AT, Yakobian N, Astourian M, et al. Inhibition of IL-17A Protects against Thyroid Immune-Related Adverse Events while Preserving Checkpoint Inhibitor Antitumor Efficacy. J Immunol 2022;209:696–709. 10.4049/jimmunol.2200244.

25 Axelrod ML, Meijers WC, Screever EM, Qin J, Carroll MG, Sun X, et al. T cells specific for α-myosin drive immunotherapy-related myocarditis. Nature 2022. 10.1038/s41586-022-05432-3.

26 Kim ST, Chu Y, Misoi M, Suarez-Almazor ME, Tayar JH, Lu H, et al. Distinct molecular and immune hallmarks of inflammatory arthritis induced by immune checkpoint inhibitors for cancer therapy. Nat Commun 2022;13:1–19. 10.1038/s41467-022-29539-3.

27 Sasson SC, Slevin SM, Cheung VTF, Nassiri I, Olsson-Brown A, Fryer E, et al. Interferon-Gamma–Producing CD8+ Tissue Resident Memory T Cells Are a Targetable Hallmark of Immune Checkpoint Inhibitor–Colitis. Gastroenterology 2021;161:1229–1244.e9. 10.1053/j.gastro.2021.06.025.

28 Luoma AM, Suo S, Williams HL, Sharova T, Sullivan K, Manos M, et al. Molecular Pathways of Colon Inflammation Induced by Cancer Immunotherapy. Cell 2020;182:655–671.e22. 10.1016/j.cell.2020.06.001.

29 Hu H, Zakharov PN, Peterson OJ, Unanue ER. Cytocidal macrophages in symbiosis with CD4 and CD8 T cells cause acute diabetes following checkpoint blockade of PD-1 in NOD mice. Proc Natl Acad Sci U S A 2020;117:31319–30. 10.1073/pnas.2019743117.

30 Collier JL, Pauken KE, Lee CAA, Patterson DG, Markson SC, Conway TS, et al. Single-cell profiling reveals unique features of diabetogenic T cells in anti-PD-1-induced type 1 diabetes mice. Journal of Experimental Medicine 2023;220:. 10.1084/jem.20221920.

31 Perdigoto AL, Deng S, Du KC, Kuchroo M, Burkhardt DB, Tong A, et al. Immune cells and their inflammatory mediators modify β cells and cause checkpoint inhibitor–induced diabetes. JCI Insight 2022;7:. 10.1172/jci.insight.156330.

32 Kao CJ, Charmsaz S, Alden SL, Brancati M, Li HL, Balaji A, et al. Immune-related events in individuals with solid tumors on immunotherapy associate with Th17 and Th2 signatures. J Clin Invest 2024;134:. 10.1172/JCI176567.

33 Bukhari S, Henick BS, Winchester RJ, Lerrer S, Adam K, Gartshteyn Y, et al. Single-cell RNA sequencing reveals distinct T cell populations in immune-related adverse events of checkpoint inhibitors. Cell Rep Med 2023;4:100868. 10.1016/j.xcrm.2022.100868.

34 von Euw E, Chodon T, Attar N, Jalil J, Koya RC, Comin-Anduix B, et al. CTLA4 blockade increases Th17 cells in patients with metastatic melanoma. J Transl Med 2009;7:1–13. 10.1186/1479-5876-7-35.

35 Ayers M, Lunceford J, Nebozhyn M, Murphy E, Loboda A, Kaufman DR, et al. IFN-γ-related mRNA profile predicts clinical response to PD-1 blockade. Journal of Clinical Investigation 2017;127:2930–40. 10.1172/JCI91190.

36 Seyedsadr M, Bang MF, McCarthy EC, Zhang S, Chen HC, Mohebbi M, et al. A pathologically expanded, clonal lineage of IL-21-producing CD4+ T cells drives inflammatory neuropathy. Journal of Clinical Investigation 2024;134:. 10.1172/JCI178602.

37 Choi J-Y, Seth A, Kashgarian M, Terrillon S, Fung E, Huang L, et al. Disruption of Pathogenic Cellular Networks by IL-21 Blockade Leads to Disease Amelioration in Murine Lupus. J Immunol 2017;198:2578–88. 10.4049/jimmunol.1601687.

38 Dong X, Antao OQ, Song W, Sanchez GM, Zembrzuski K, Koumpouras F, et al. Type I Interferon-Activated STAT4 Regulation of Follicular Helper T Cell-Dependent Cytokine and Immunoglobulin Production in Lupus. Arthritis Rheumatol 2021;73:478–89. 10.1002/art.41532.

39 Reinhardt RL, Liang H-E, Locksley RM. Cytokine-secreting follicular T cells shape the antibody repertoire. Nat Immunol 2009;10:385–93. 10.1038/ni.1715.

40 Pauken KE, Jenkins MK, Azuma M, Fife BT. PD-1, but not PD-L1, expressed by islet-Reactive CD4+ T cells suppresses infiltration of the pancreas during type 1 diabetes. Diabetes 2013;62:2859–69. 10.2337/db12-1475.

41 Spolski R, Leonard WJ. Interleukin-21: A double-edged sword with therapeutic potential. Nat Rev Drug Discov 2014;13:379–95. 10.1038/nrd4296.

42 Leonard WJ, Wan CK. IL-21 Signaling in Immunity. F1000Res 2016;5:1–10. 10.12688/f1000research.7634.1.

43 Spolski R, Kashyap M, Robinson C, Yu Z, Leonard WJ. IL-21 signaling is critical for the development of type I diabetes in the NOD mouse. Proc Natl Acad Sci U S A 2008;105:14028–33. 10.1073/pnas.0804358105.

44 Ciecko AE, Wang Y, Harleston S, Drewek A, Serreze D V., Geurts AM, et al. Heterogeneity of Islet-Infiltrating IL-21+ CD4 T Cells in a Mouse Model of Type 1 Diabetes. The Journal of Immunology 2023;210:935–46. 10.4049/jimmunol.2200712.

45 Wang S-P, Iwata S, Nakayamada S, Sakata K, Yamaoka K, Tanaka Y. Tofacitinib, a JAK inhibitor, inhibits human B cell activation in vitro. Ann Rheum Dis 2014:2213–5. 10.1136/annrheumdis-2014-205615.

46 Katarzyna PB, Wiktor S, Ewa D, Piotr L. Current treatment of systemic lupus erythematosus: a clinician’s perspective. Rheumatol Int 2023;43:1395–407. 10.1007/s00296-023-05306-5.

47 Guo J, Zhang H, Lin W, Lu L, Su J, Chen X. Signaling pathways and targeted therapies for psoriasis. Signal Transduct Target Ther 2023;8:. 10.1038/s41392-023-01655-6.

48 Cai Z, Wang S, Li J. Treatment of Inflammatory Bowel Disease: A Comprehensive Review. Front Med (Lausanne) 2021;8:1–24. 10.3389/fmed.2021.765474.

49 Platanias LC. Mechanisms of type-I- and type-II-interferon-mediated signalling. Nat Rev Immunol 2005;5:375–86. 10.1038/nri1604.

50 Fife BT, Guleria I, Bupp MG, Eagar TN, Tang Q, Bour-Jordan H, et al. Insulin-induced remission in new-onset NOD mice is maintained by the PD-1-PD-L1 pathway. Journal of Experimental Medicine 2006;203:2737–47. 10.1084/jem.20061577.

51 Ansari MJI, Salama AD, Chitnis T, Smith RN, Yagita H, Akiba H, et al. The programmed death-1 (PD-1) pathway regulates autoimmune diabetes in nonobese diabetic (NOD) mice. Journal of Experimental Medicine 2003;198:63–9. 10.1084/jem.20022125.

52 Paterson AM, Brown KE, Keir ME, Vanguri VK, Riella L V, Chandraker A, et al. The programmed death-1 ligand 1:B7-1 pathway restrains diabetogenic effector T cells in vivo. J Immunol 2011;187:1097–105. 10.4049/jimmunol.1003496.

53 Wang Z, Zhao M, Yin J, Liu L, Hu L, Huang Y, et al. E4BP4-mediated inhibition of T follicular helper cell differentiation is compromised in autoimmune diseases. Journal of Clinical Investigation 2020;130:3717–33. 10.1172/JCI129018.

54 Marsiglio J, McPherson JP, Kovacsovics-Bankowski M, Jeter J, Vaklavas C, Swami U, et al. A single center case series of immune checkpoint inhibitor-induced type 1 diabetes mellitus, patterns of disease onset and long-term clinical outcome. Front Immunol 2023;14:1–11. 10.3389/fimmu.2023.1229823.

55 Ge T, Phung AL, Jhala G, Trivedi P, Principe N, De George DJ, et al. Diabetes induced by checkpoint inhibition in nonobese diabetic mice can be prevented or reversed by a JAK1/JAK2 inhibitor. Clin Transl Immunology 2022;11:1–16. 10.1002/cti2.1425.

56 Mathew D, Marmarelis ME, Foley C, Bauml JM, Ye D, Ghinnagow R, et al. Combined JAK inhibition and PD-1 immunotherapy for non–small cell lung cancer patients. Science (1979) 2024;384:eadf1329. 10.1126/science.adf1329.

57 Zak J, Pratumchai I, Marro BS, Marquardt KL, Zavareh RB, Lairson LL, et al. JAK inhibition enhances checkpoint blockade immunotherapy in patients with Hodgkin lymphoma. Science 2024;384:eade8520. 10.1126/science.ade8520.

58 Senkevitch E, Li W, Hixon JA, Andrews C, Cramer SD, Pauly GT, et al. Inhibiting Janus Kinase 1 and BCL-2 to treat T cell acute lymphoblastic leukemia with IL7-Rα mutations. Oncotarget 2018;9:22605–17. 10.18632/oncotarget.25194.

59 Villarreal D, Pradhan G, Wu C-S, Allred CD, Guo S, Sun Y. A Simple High Efficiency Protocol for Pancreatic Islet Isolation from Mice. J Vis Exp 2019. 10.3791/57048.

60 Lo B, Swafford ADE, Shafer-Weaver KA, Jerome LF, Rakhlin L, Mathern DR, et al. Antibodies against insulin measured by electrochemiluminescence predicts insulitis severity and disease onset in non-obese diabetic mice and can distinguish human type 1 diabetes status. J Transl Med 2011;9:1–16. 10.1186/1479-5876-9-203.

61 You S, Chen C, Lee W-H, Wu C-H, Judkowski V, Pinilla C, et al. Detection and Characterization of T Cells Specific for BDC2.5 T Cell-Stimulating Peptides. The Journal of Immunology 2003;170:4011–20. 10.4049/jimmunol.170.8.4011.

