## Supplemental Tables and Figures for "Polyfunctional IL-21^+^ IFNγ^+^ T follicular helper cells contribute to checkpoint inhibitor diabetes mellitus and can be targeted by JAK inhibitor therapy"

| | Age (years) at start of ICI | Sex | Immunotherapy | Cancer type | IrAEs |
|---|---|---|---|---|---|
| **Individuals with No IRAEs (Fig. 1)** | 47 | Male | Pembrolizumab | Metastatic Melanoma | No significant IrAE |
| | 43 | Female | Pembrolizumab | Head and Neck SCC | No significant IrAE |
| | 47 | Female | Pembrolizumab | Diffuse type Gastric Carcinoma | No significant IrAE |
| | 55 | Male | Pembrolizumab | Head and Neck SCC | No significant IrAE |
| | 75 | Male | Atezolizumab | Pancreatic Cancer | No significant IrAE |
| | **Age (years) at IrAE** | **Sex** | **Immunotherapy** | **Cancer type** | **IrAEs** |
| **Individuals with ICI-T1DM (Fig. 1)** | 64 | Male | Nivolumab | Metastatic Melanoma | ICI-T1DM |
| | 53 | Male | Pembrolizumab | Metastatic Melanoma | ICI-T1DM, ICI-Thyroiditis, ICI-Hypophysitis |
| | 60 | Male | Nivolumab | Metastatic Melanoma | ICI-T1DM |
| | 56 | Female | Nivolumab | Mucosal Melanoma of Vagina | ICI-T1DM, ICI-Pneumonitis, ICI-Colitis |
| | 59 | Female | Nivolumab | Melanoma | ICI-T1DM, ICI-thyroiditis, ICI-Sjogrens |
| | **Age (years) at IrAE** | **Sex** | **Immunotherapy** | **Cancer Type** | **IrAEs** |
| **ICI-treated patients (Fig. 5)** | 65 | Male | Nivolumab | Metastatic Melanoma | ICI-T1DM, ICI-Thyroiditis |
| | 69 | Male | Pembrolizumab | Metastatic Melanoma | ICI-T1DM, ICI-Thyroiditis, ICI-Colitis, ICI-Arthritis |
| | 71 | Female | Pembrolizumab | Triple Negative Breast Cancer | ICI-Thyroiditis, ICI-Hypophysitis |
| | 34 | Female | Nivolumab | Metastatic Melanoma | ICI-Hypophysitis, ICI-Hepatitis |
| | 70 | Male | Pembrolizumab | Urothelial Cancer | ICI-Thyroiditis, ICI-Arthritis, ICI-Colitis |

**Supplemental Table 1. Demographic and clinical data for patient specimens.** ICI, Immune checkpoint inhibitor; IrAE, Immune related Adverse Event; T1DM, type 1 diabetes mellitus; SCC, squamous cell carcinoma.

**Mouse antibodies**

| Target | Fluorescence | Clone | Vendor |
|---|---|---|---|
| CD4 | e450 | GK1.5 | Invitrogen |
| PD-1 | APC | J43 | Invitrogen |
| ICOS | PerCPCy5.5 | C398.4A | Biolegend |
| CXCR5 | FITC | L138D7 | Biolegend |
| BCL6 | APC-Cy7 | K112-91 | BD Biosciences |
| cMAF | PE-Cy7 | sym0F1 | Invitrogen |
| IL-21 | PE | mhalx21 | Invitrogen |
| IFNy | APC | XMG1.2 | Biolegend |
| pSTAT3 | unconjugated | D3A7 | Cell Signaling |
| PD-1 | PE-Cy7 | J43 | Invitrogen |
| CD4 | PE | RM4-5 | Biolegend |
| CD8 | APC | 53-6.7 | Biolegend |

**Human anitbodies**

| Target | Fluorescence | Clone | Vendor |
|---|---|---|---|
| CD4 | PE | RPA-T4 | Biolegend |
| CXCR5 | AF488 | J252D4 | Biolegend |
| ICOS | PE-Cy7 | ISA-3 | Invitrogen |
| PD-1 | e450 | MIH4 | Invitrogen |
| BCL6 | APC-Cy7 | K112-91 | BD Biosciences |

**Suplemental Table 2. Flow cytometry antibodies.**

### Supplemental Figures

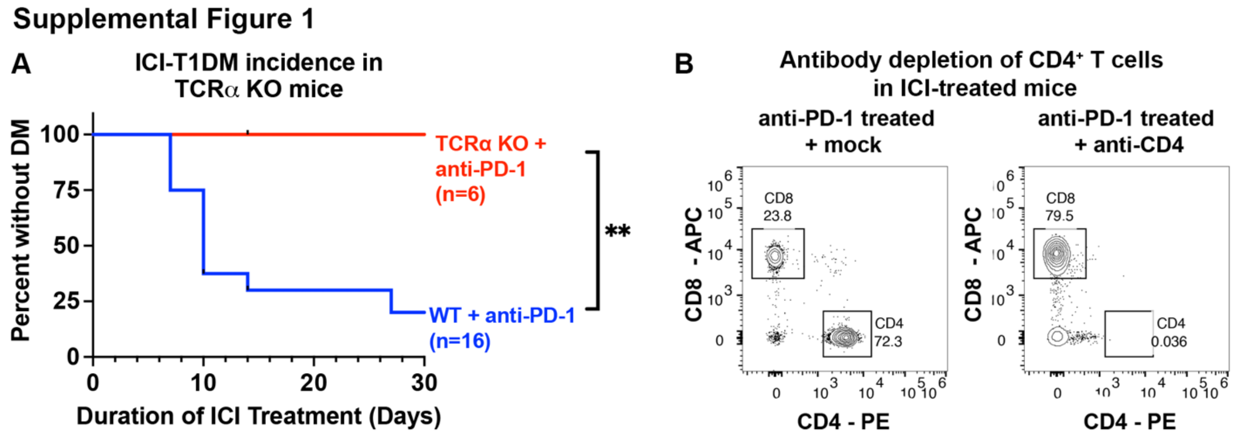


**Supplemental Figure 1. CD4$^{+}$ and CD8$^{+}$ T cells are required for the development of immune checkpoint inhibitor autoimmune diabetes in NOD mice.**

A. Incidence of autoimmune diabetes mellitus (DM) in NOD wildtype (WT) or NOD.TCRα$^{-/-}$ (TCRα KO) mice treated with anti-PD-1 or isotype control. Comparison by Log Rank test, **p<0.01.
B. Representative flow cytometry plots of splenocytes from NOD mice treated anti-PD1 demonstrating CD4$^{+}$ T cell depletion with anti-CD4 antibody.

**Supplemental Figure 2**

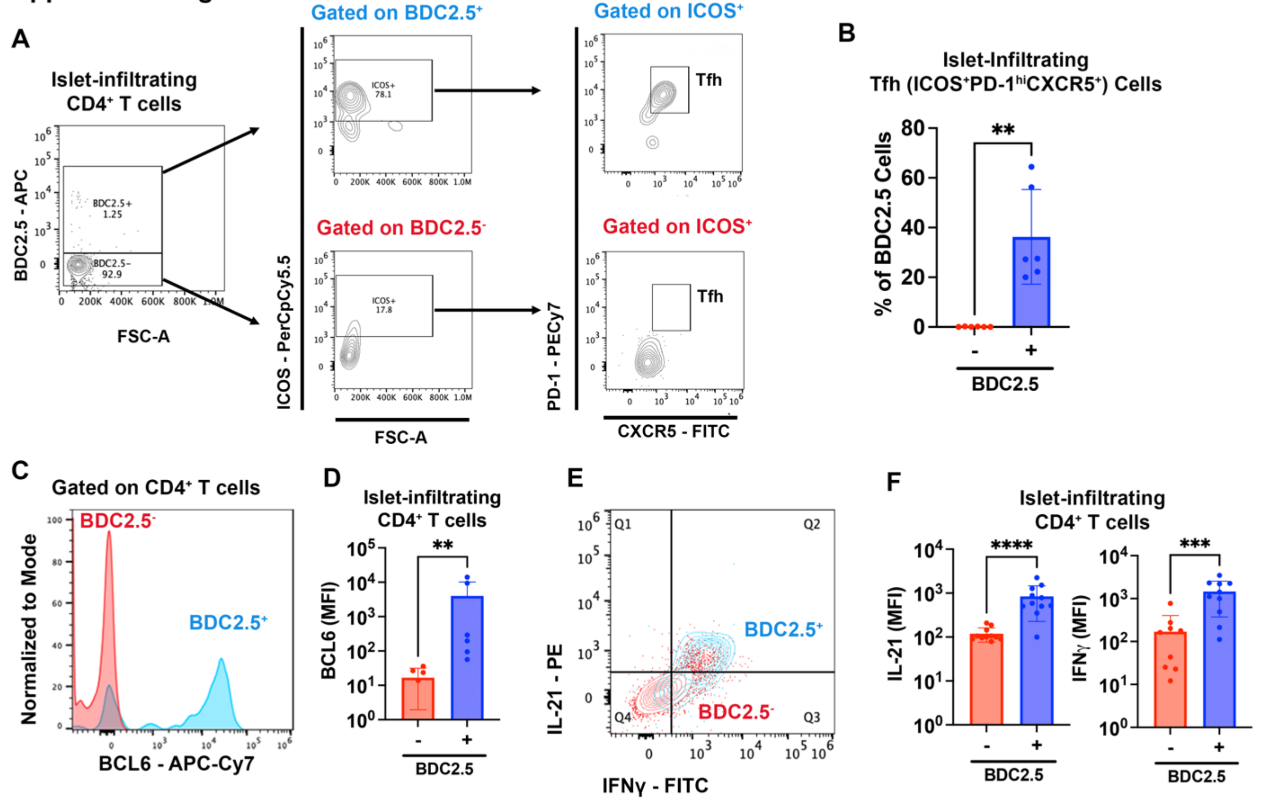


**Supplemental Figure 2. BDC2.5 tetramer staining reveals antigen-specific Tfh CD4$^+$ T cells within pancreatic islets of ICI-treated mice.**

A. Representative flow cytometry plots of BDC2.5$^+$ tetramer CD4$^+$ T cells within the pancreatic islets of anti-PD-1 treated mice showing gating for putative T follicular helper (Tfh) cell surface markers.
B. Frequency of Tfh (ICOS$^+$ PD-1$^{hi}$ CXCR5$^+$) cells among BDC2.5$^+$ versus BDC2.5$^-$ CD4$^+$ T cells within pancreatic islets of anti-PD-1 treated mice.
C. Representative histogram of Bcl6 staining among BDC2.5$^+$ versus BDC2.5$^-$ CD4$^+$ T cells within pancreatic islets.
D. Comparison of Bcl6 expression among BDC2.5$^+$ versus BDC2.5$^-$ CD4$^+$ T cells within pancreatic islets in anti-PD-1 treated mice.
E. Representative flow plot of IL-21 and IFNγ expression of BDC2.5$^+$ and CDC2.5$^-$ CD44$^+$ CD4$^+$ T cells within pancreatic islets from anti-PD-1 treated mice.
F. Comparison of IL-21 and IFNγ expression among BDC2.5$^+$ versus BDC2.5$^-$ CD4$^+$ T cells within pancreatic islets in anti-PD-1 treated mice.

Absolute cell counts and frequencies of islet-infiltrating cell types were determined by flow cytometry. Each point represents data from one animal and data are presented as mean±SD. Comparisons by Mann-Whitney test (B, D, F). **p<0.01; ***p<0.001, ****p<0.0001. FSC, forward scatter; PerCP-Cy5.5, peridinin chlorophyll protein-cyanine

5; PECy-7, phycoerythrin-cyanine 7; FITC, fluorescein isothiocyanate; APC, allophycocyanin; PE, phycoerythrin; FITC, fluorescein isothiocyanate; APC-Cy7, allophycocyanin-cyanine 7.

Supplemental Figure 3

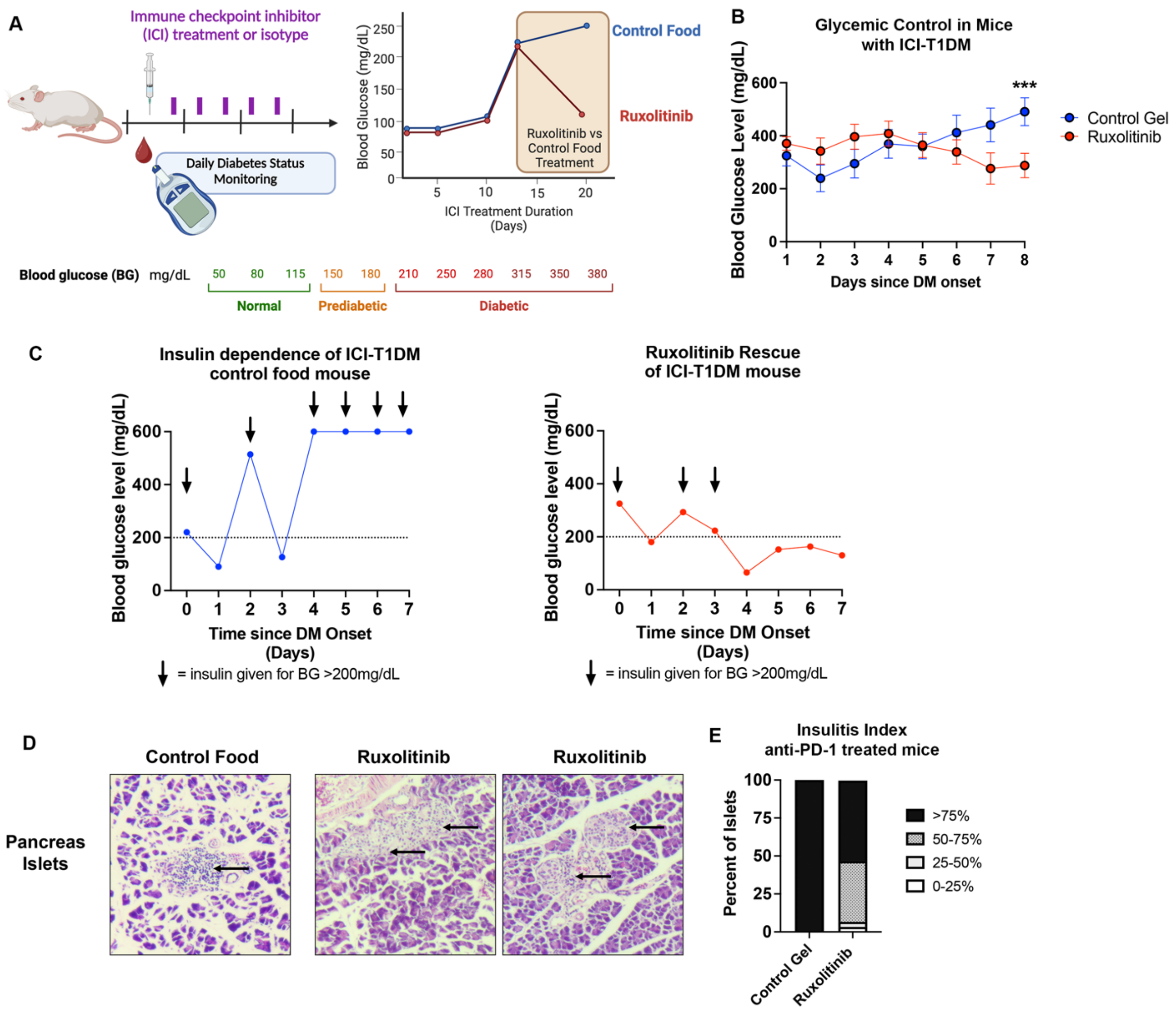


**Supplemental Figure 3. Ruxolitinib improves glycemic control and reduces insulitis in ICI-T1DM even after onset of overt disease.**

A. Schematic of ruxolitinib reversal trial in mice with ICI-T1DM. Mice were treated with anti-PD-1 twice weekly and monitored for development of hyperglycemia (blood glucose >200mg/dL) daily. Once mice developed hyperglycemia, they were randomized to treatment with JAKi ruxolitinib or control food.
B. Comparison of glycemic control mice with ICI-T1DM treated with ruxolitinib or control food gel. Data are presented as mean±SD. Comparisons by unpaired Student's t test on day eight. Data combined from three independent experiments. ***$p<0.001$.
C. Blood glucose and insulin requirements in a mouse with ICI-T1DM given control food (*left*) compared to a mouse given ruxolitinib (*right*) after onset of hyperglycemia.

D. Representative hematoxylin and eosin staining of pancreas histology showing islets of Langerhans.
E. Insulitis index of pancreas islet histology of mice with ICI-T1DM given ruxolitinib or control food gel after onset of overt diabetes.

### Supplemental Figure 4

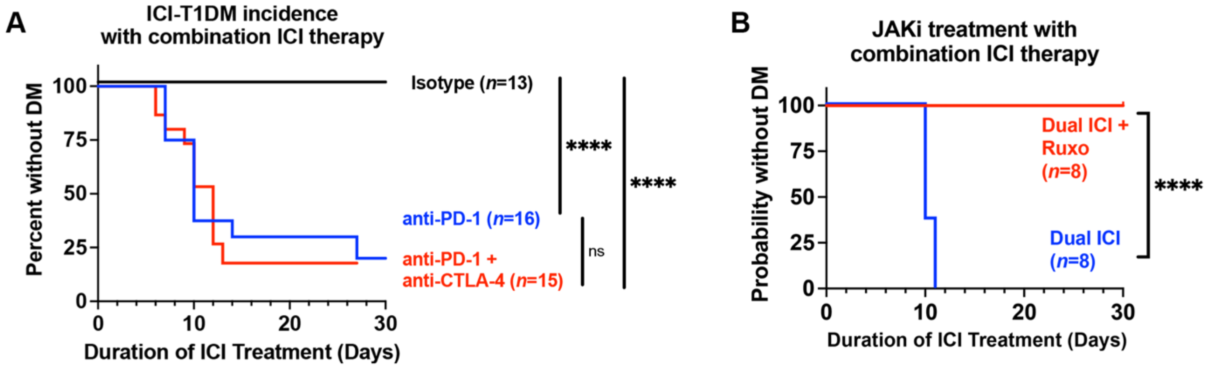


**Supplemental Figure 4. Ruxolitinib prevents immune checkpoint inhibitor (ICI) autoimmune diabetes mellitus during combination ICI therapy.**

A. Incidence of autoimmune diabetes mellitus (DM) in NOD mice treated with isotype, anti-PD-1 (data reshown from Fig. 2B), and combination anti-PD-1 + anti-CTLA-4.
B. ICI-T1DM incidence in NOD mice treated with isotype or Dual ICI (anti-PD-1 + anti-CTLA-4) in combination with ruxolitinib (1g/kg daily) or control food.
   Comparisons by Log-Rank test (A, B). ****p<0.0001.
